# Cell culture-based production of defective interfering influenza A virus particles in perfusion mode using an alternating tangential flow filtration system

**DOI:** 10.1101/2021.06.07.446880

**Authors:** Marc D. Hein, Anshika Chawla, Maurizio Cattaneo, Sascha Y. Kupke, Yvonne Genzel, Udo Reichl

**Affiliations:** Otto-von-Guericke-University Magdeburg, Chair of Bioprocess Engineering, Magdeburg, Germany; Max Planck Institute for Dynamics of Complex Technical Systems, Bioprocess Engineering, Magdeburg, Germany; Martin Luther University Halle-Wittenberg, Institute of Pharmacy, Halle (Saale), Germany; Artemis Biosystems, Cambridge, Massachusetts, USA

**Keywords:** influenza A virus, antiviral, defective interfering particles (DIPs), cell culture-based production, perfusion cultivation, alternating tangential flow filtration (ATF), bioreactor

## Abstract

Respiratory diseases including influenza A virus (IAV) infections represent a major threat to human health. While the development of a vaccine requires a lot of time, a fast countermeasure could be the use of defective interfering particles (DIPs) for antiviral therapy. IAV DIPs are usually characterized by a large internal deletion in one viral RNA segment. Consequentially, DIPs can only propagate in presence of infectious standard viruses (STVs), compensating the missing gene function. Here, they interfere with and suppress the STV replication and might act “universally” against many IAV subtypes. We recently reported a production system for purely clonal DIPs utilizing genetically modified cells. In the present study, we established an automated perfusion process for production of a DIP, called DI244, using an alternating tangential flow filtration (ATF) system for cell retention. Viable cell concentrations and DIP titers more than 10-times higher than for a previously reported batch cultivation were observed. Further, we investigated a novel tubular cell retention device for its potential for continuous virus harvesting into the permeate. Very comparable performances to typically used hollow fiber membranes were found during the cell growth phase. During the virus replication phase the tubular membrane, in contrast to the hollow fiber membrane, allowed 100% of the produced virus particles to pass through. To our knowledge, this is the first time a continuous virus harvest was shown for a membrane-based perfusion process. Overall, the process established offers interesting possibilities for advanced process integration strategies for next-generation virus particle and virus vector manufacturing.

**Key points:**

- An automated perfusion process for production of IAV DIPs was established
- DIP titers of 7.40E+9 plaque forming units per mL were reached
- A novel tubular cell retention device enabled continuous virus harvesting

## Introduction

Infections with influenza A virus (IAV; list of abbreviations, Tab. 1) result worldwide in up to 650,000 deaths annually (Iuliano et al. 2018). The potential of a pandemic is a constant threat, as IAV pandemics have led to millions of deaths in the past and highly infectious variants of this RNA virus can emerge at any time (Johnson and Mueller 2002; Taubenberger et al. 2000). In case of such a pandemic situation, the use of antivirals could be crucial as a first line of defense to avoid a high number of deaths until a vaccine is available. Additionally, antivirals could be used to supplement vaccination during annual epidemics. Currently used antivirals against IAV include oseltamivir and zanamivir (Colman 2009; Oxford 2007; Smith et al. 2006). However, resistances of some IAV strains against mentioned antivirals have already been reported (Han et al. 2018; Lackenby et al. 2018) clearly demonstrating the need for development of novel antiviral drugs, preferably with a broad efficacy spectrum.

**Table 1.**
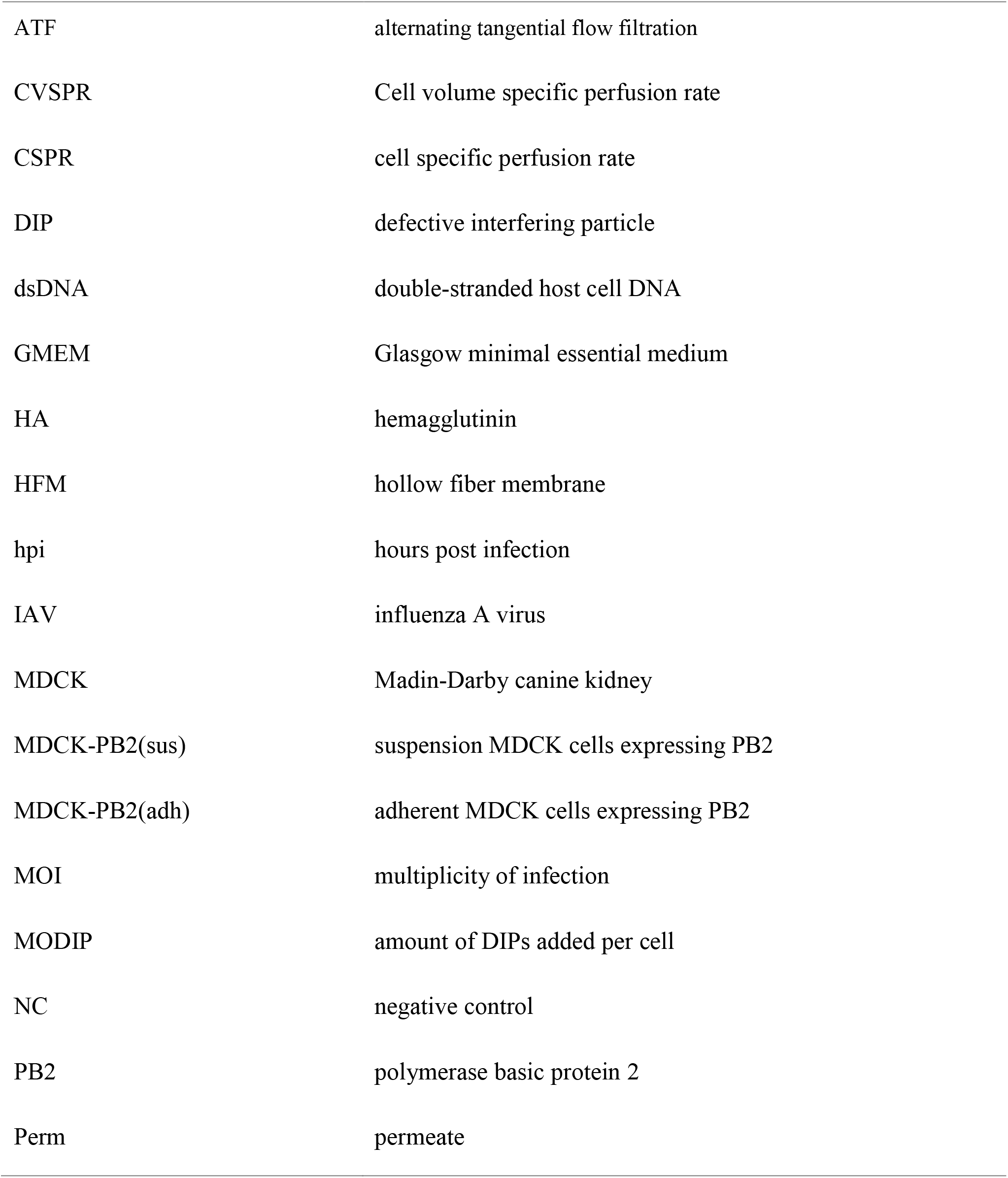
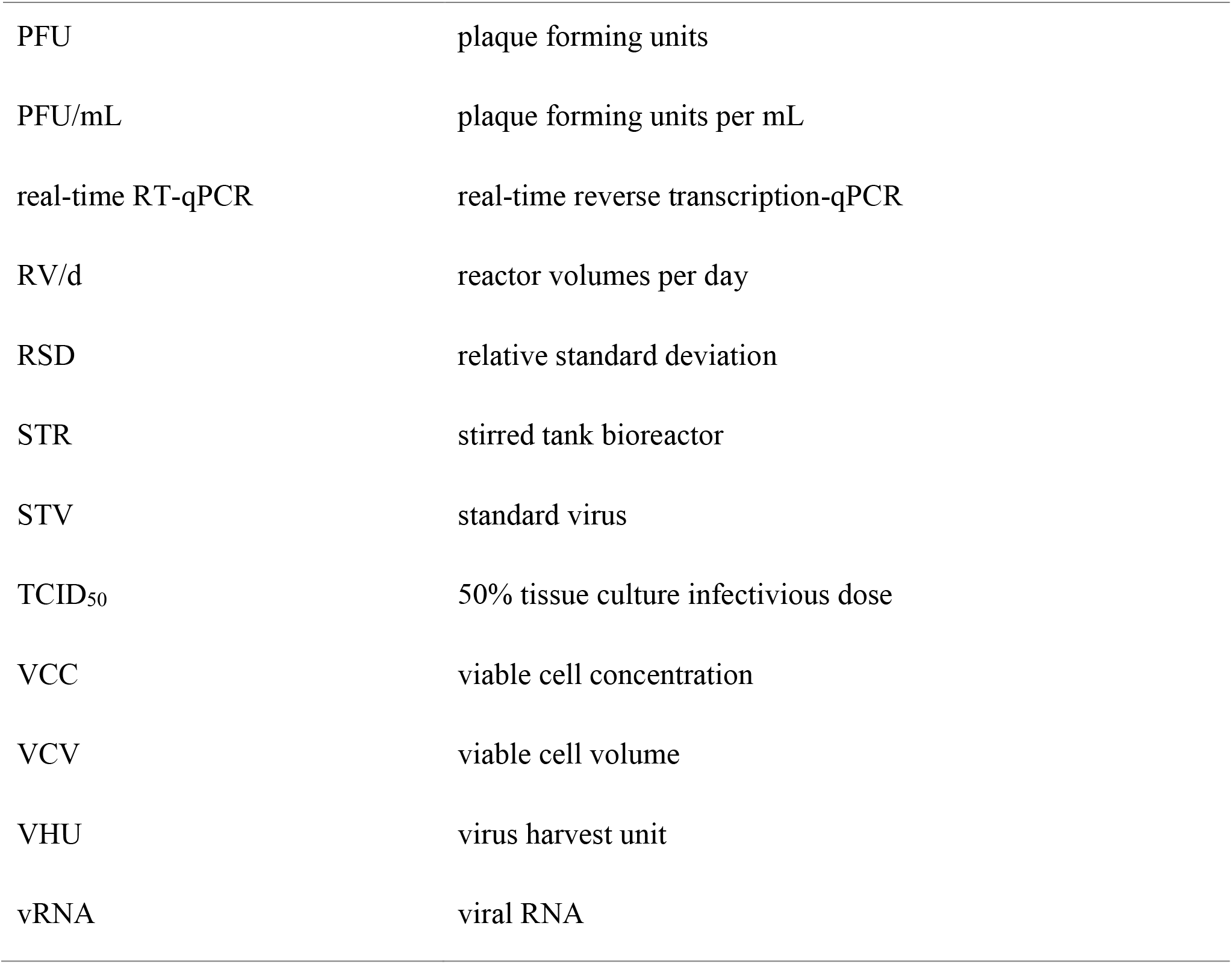
List of Abbreviations.

|  |  |
| --- | --- |
| ATF | alternating tangential flow filtration |
| CVSPR | Cell volume specific perfusion rate |
| CSPR | cell specific perfusion rate |
| DIP | defective interfering particle |
| dsDNA | double-stranded host cell DNA |
| GMEM | Glasgow minimal essential medium |
| HA | hemagglutinin |
| HFM | hollow fiber membrane |
| hpi | hours post infection |
| IAV | influenza A virus |
| MDCK | Madin-Darby canine kidney |
| MDCK-PB2(sus) | suspension MDCK cells expressing PB2 |
| MDCK-PB2(adh) | adherent MDCK cells expressing PB2 |
| MOI | multiplicity of infection |
| MODIP | amount of DIPs added per cell |
| NC | negative control |
| PB2 | polymerase basic protein 2 |
| Perm | permeate |
| PFU | plaque forming units |
| PFU/mL | plaque forming units per mL |
| real-time RT-qPCR | real-time reverse transcription-qPCR |
| RV/d | reactor volumes per day |
| RSD | relative standard deviation |
| STR | stirred tank bioreactor |
| STV | standard virus |
| TCID<sub>50</sub> | 50% tissue culture infectivious dose |
| VCC | viable cell concentration |
| VCV | viable cell volume |
| VHU | virus harvest unit |
| vRNA | viral RNA |

Discussion on such possible alternative antivirals was started when defective interfering particles (DIPs) were proposed for the use in antiviral therapy (Dimmock et al. 1986; Dimmock et al. 2008). DIPs are virus mutants occurring naturally during the error-prone replication of the viral genome (Lazzarini et al. 1981; Perrault 1981). The majority of IAV DIPs is characterized by a large deletion in the open reading frame of one of their eight viral RNA (vRNA) segments (Nayak et al. 1985). Thus, DIPs lack the genetic information necessary for the synthesis of the corresponding full-length protein (Huang and Baltimore 1970). Consequentially, DIPs are unable to replicate, unless the missing protein is provided, for example by a co-infection with the standard virus (STV). In such a co-infection, DI vRNAs seem to replicate faster than the full-length equivalent and therefore outcompete the STV vRNA for limited cellular resources (Dimmock and Easton 2014; Laske et al. 2016; Nayak et al. 1985). In addition, other mechanisms might also play a role in suppressing STV replication and spreading (Easton et al. 2011; Scott et al. 2011). Overall, this results in a drastic decrease in released infectious virus particles, as mainly non-infectious DIPs are produced (Frensing 2015; Tapia et al. 2019; Von Magnus 1951). The DIP-induced inhibition of STV replication is causative for their antiviral potential, previously demonstrated in mice (Dimmock et al. 2008). Additionally, the antiviral effect of IAV DIPs against influenza B, pneumovirus and SARS-CoV2 was shown, likely mediated by an unspecific stimulation of the innate immune response (Easton et al. 2011; Rand et al. 2021; Scott et al. 2011).

Early methods to produce IAV DIPs focused on egg-based manufacturing with STV co-infection (Dimmock et al. 2008). Unfortunately, this resulted in a relatively high variability of the produced material (contamination with multiple other DIPs). Furthermore, it required the inactivation of the infectious STV after harvesting, for example by UV irradiation. This UV irradiation also inactivated parts of the produced DIPs and therefore decreased the antiviral potency of final preparations used in animal trials (Hein et al. 2021b). To overcome this limitation, a method for production of purely clonal DIP populations, devoid of STV and other DIPs, was developed (Bdeir et al. 2019). Here, adherent Madin-Darby canine kidney (MDCK) cells were genetically modified to express the polymerase basic protein 2 (PB2) encoded by IAV segment 1 (Seg1) vRNA. This allowed propagation of IAV DIPs with a deletion in Seg1. We recently reported the production of a prototypic Seg1 DIP, called DI244, utilizing a suspension MDCK cell line expressing PB2 (MDCK-PB2(sus)) and the evaluation of the produced DI244 in animal experiments (Hein et al. 2021a). Here, all mice infected with an otherwise lethal dose of IAV survived the infection when treated with DI244. These experiments already clearly demonstrated the antiviral potential of these purely clonal DIPs. However, the DI244 material evaluated in this study was produced in shake flasks only and large DIP quantities were required. Consequentially, it was yet to be demonstrated that a scalable high-yield process could be implemented in a stirred tank bioreactor (STR).

In the present study, we therefore propose process intensification strategies to further improve the cell culture-based production of IAV DIPs. First, a perfusion process to grow MDCK-PB2(sus) cells to high cell concentrations was established that utilized an alternating tangential flow filtration system (ATF) with a standard hollow fiber membrane (HFM, pore size of 0.2 μm). The perfusion rate was adjusted manually every 12 hours. In a next step, the perfusion rate was controlled based on cell concentration measurements using a capacitance probe. Sufficient substrate and low waste product levels were maintained and a cell concentration of 28.4E+06 cells/mL was reached, resulting in a DI244 titer of 7.40E+9 plaque forming units per mL (PFU/mL). Lastly, a tubular membrane called virus harvest unit (VHU) was used for continuous virus particle harvesting into the permeate during the perfusion process.

## Materials and Methods

### Cells and viruses

MDCK-PB2(sus) cells were used for production of propagation incompetent DI244 as described previously (Hein et al. 2021a). Briefly, MDCK cells adapted to growth in chemically defined Xeno™ medium (Bissinger et al. 2019) were genetically modified by retroviral transduction to express the viral PB2, encoded by IAV Seg1 vRNA. The PB2 expression allowed propagation of pure DI244 particles, harboring a deletion in Seg1. Additionally, an adherent MDCK cell line expressing PB2 (Bdeir et al. 2019), hereafter MDCK-PB2(adh), was used for DI244 quantification (see “virus quantification assays”).

MDCK-PB2(sus) cells were maintained in a shake flask with 40 mL working volume (125 mL baffled polycarbonate Erlenmeyer flask, Thermo Fisher Scientific, 4116-0125) at 37°C, 5% CO_2_ and 185 rpm (Multitron Pro, Infors HT; 50 mm shaking orbit). Cells were passaged every 2–3 days. For DI244 production, MDCK-PB2(sus) cells were grown in a STR in perfusion mode (see “perfusion cultivation of MDCK-PB2(sus) cells”). The parental adherent MDCK cells (ECACC, No. 84121903) and the genetically modified MDCK-PB2(adh) were cultivated in Glasgow minimum essential medium (GMEM) containing 1% peptone and 10% fetal bovine serum at 37°C and 5% CO_2_. Viable cell concentration (VCC), cell viability, and cell diameter of suspension and adherent cells were determined using an automated cell counter (Vi-CELL XR, Beckman Coulter, 731050).

A pure DI244 seed virus was generated using an eight-plasmid DNA transfection system, as described previously (Bdeir et al. 2019). For infection of MDCK-PB2(sus) cells, the amount of DIPs added per cell (multiplicity of DIP (MODIP)) was calculated based on the DI244 titer of the used seed virus (8.40E+07 PFU/mL). For STV infections of adherent MDCK cells (see “interference assay”) the influenza virus A/Puerto Rico/8/34 (provided by the Robert Koch Institute, Germany; Amp. 3138) was used. The multiplicity of infection (MOI) was calculated based on the tissue culture infectious dose (TCID_50_) titer (Genzel and Reichl 2007) of the MDCK adapted seed virus (1.1E+9 TCID_50_/mL).

### Perfusion cultivation of MDCK-PB2(sus) cells

The implementation of a perfusion process utilizing the Xeno™ medium but a different MDCK cell line was reported previously (Wu et al. 2021). In the present study, the STR (DASGIP^®^ Parallel Bioreactor System, Eppendorf AG, 76DG04CCBB, working volume 700 mL) used for DIP production was equipped with one inclined blade impeller (three blades, 30° angle, 50 mm diameter) and two spargers (one macro- and one micro-sparger). The micro-sparger was only used for additional gassing, when the VCC increased to concentrations where gassing with the macro-sparger alone was not sufficient anymore to maintain a pO_2_ above 40%. Before inoculation, cells grown in shake flasks (as described before) were centrifuged (300×g, 5 min, room temperature) and the medium exchanged. The STR was inoculated with a VCC of 1.0E+6 cells/mL and operated at 37°C, pO_2_≥40%, pH 7.5 and 150 rpm. Before perfusion was started, the bioreactor was operated for 24 h in batch mode.

### Cell retention using an alternating tangential flow filtration system

An alternating tangential flow filtration system (ATF 2, Repligen) was used for cell retention. The flow rate of the diaphragm pump was set to 0.9 L/min, other parameters of the ATF 2 controller were kept as given by the supplier. In this study, two different membranes were investigated. One was the commonly used polyethersulfone HFM (0.2 μm pore size, 470 cm^2^ surface area, Spectrum Labs), the other one was the tubular VHU (~10 μm pore size, 60 cm^2^ surface area, Artemis Biosystems). For the removal of cell free supernatant, a cross flow through the membrane (hereafter called permeate flow) was applied by the peristaltic pump.

### Control of the perfusion rate

The feed of cultivation medium (Xeno™ medium) was controlled by a scale under the STR to maintain a constant working volume. Therefore, the feed flow rate always equaled the permeate flow rate. The permeate flow rate was increased over the process time either manually or automatically to maintain a cell specific perfusion rate (CSPR) of approximately 200 pL/cell/day. For the manually adjusted perfusion rate, a perfusion profile was calculated based on the growth rate and metabolite consumption rates determined during the exponential phase of a previous batch cultivation (Hein et al. 2021a). Here, the perfusion rate was increased stepwise every 12 hours. For the automated perfusion, a previously reported approach was used (Nikolay et al. 2018). Here, a multi-frequency capacitance probe connected to a controller (ArcView Controller 265, Hamilton) was utilized. The controller converted the measured capacity to a permittivity signal, which was correlated to the VCC in the STR (in the present study the permittivity signal was correlated to the viable cell volume (VCV) instead). The permittivity signal was forwarded to an analog 4 – 20 mA output box (Hamilton), which then again was connected to the peristaltic pump (120 U, Watson-Marlow). The desired cell volume specific perfusion rate (CVSPR) was realized by adjusting the specific cell factor stored in the ArcView controller. In this study, a cell factor of 1.18 was used to achieve a CVSPR of 0.12 pL/μm^3^/d. This would equal a CSPR of 200 pL/cell/d, when a cell is assumed to be a perfect sphere with a diameter of 15 μm. This control regime could only be used prior to virus addition as the permittivity signal was drastically disturbed during the infection phase. Therefore, the perfusion rate was kept constant after infection. The overall set up including the devices for perfusion control is shown in Fig. 1.

**Fig. 1.**
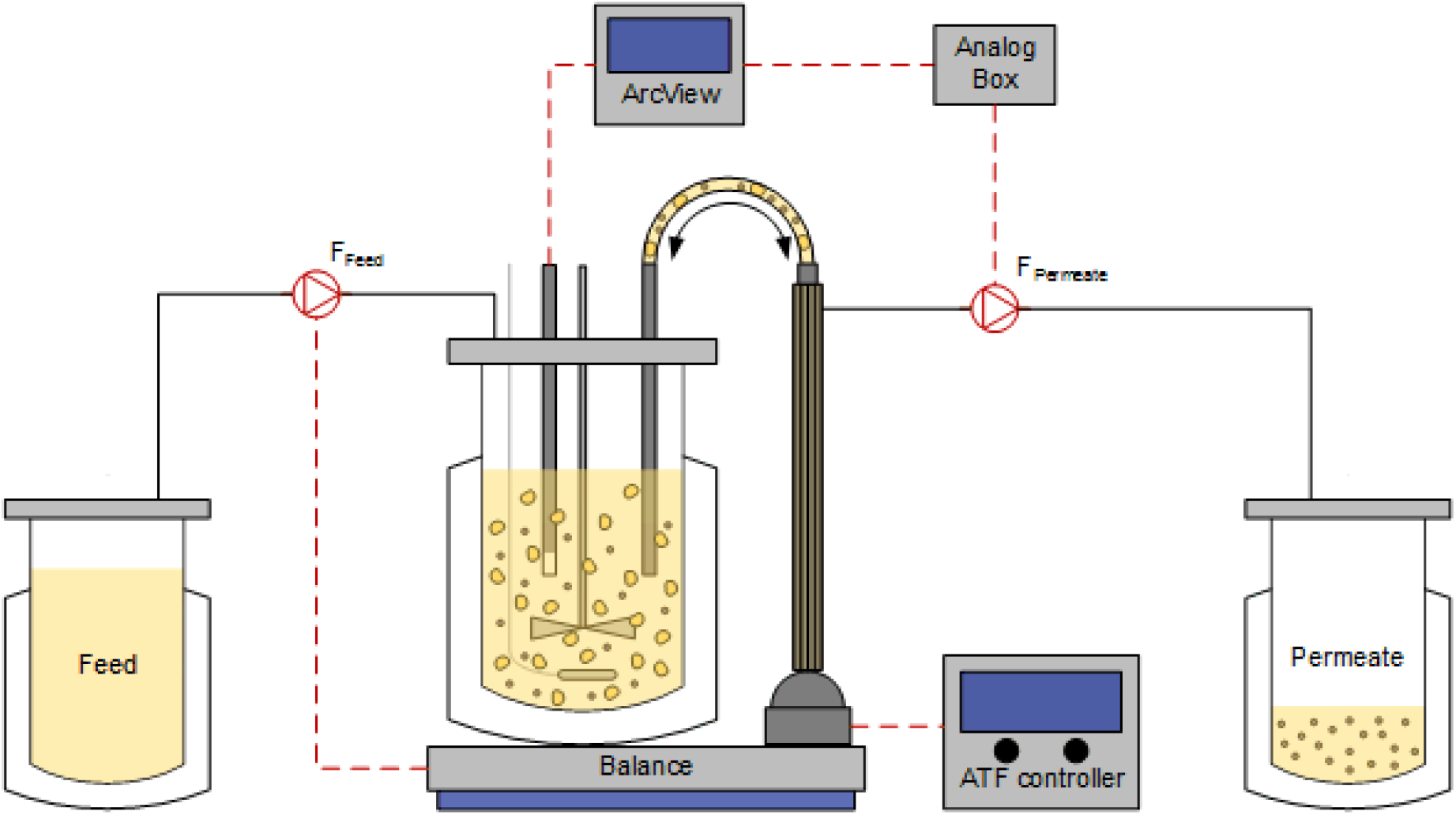
Scheme of the bioreactor setup for perfusion cultivations (according to (Nikolay et al. 2018)). For cell retention, an alternating tangential flow filtration system (ATF 2, Repligen) was utilized. A capacitance probe was used for online monitoring of the viable cell volume. The signal was converted by the ArcView and analog box to control a peristaltic pump and thus the perfusion rate. Here, a cell volume specific perfusion rate of 0.12 pL/μm^3^/d, corresponding to a cell specific perfusion rate of 200 pL/cell/d, was realized. Depending on the membrane used for cell retention, virus particles accumulate either exclusively in the cultivation vessel or additionally in the permeate. Black lines indicate tubes, red dashed lines indicate different types of signal transmission.

### Infection of cells grown in a STR

When the desired cell concentration of 20.0 – 25.0E+6 cells/mL was reached, the seed virus was added. To improve virus replication, a complete medium exchange was performed prior to infection. This was realized by drastically increasing the perfusion rate manually. When a HFM was used, the perfusion rate was increased to 100 reactor volumes per day (RV/d) for the medium exchange. In contrast, for the VHU the perfusion rate was only increased to 15 RV/d. For higher perfusion rates, damage of the VHU was observed in previous perfusion cultivations (data not shown). After the medium exchange, the cultivation temperature was set to 32°C, as this was shown to have a positive impact on virus replication (Wu et al. 2021). For infection, a pure DI244 seed virus at MODIP 1E–3 and 20 U/mL trypsin (5000 U/mL in PBS; 27250018, Gibco) were added. The perfusion rate was set to 0 RV/d for 1 h to allow for efficient virus entry into the cells. After this incubation time, perfusion was started again and the perfusion rate was kept constant at 2 – 3 RV/d. The perfusion medium used after infection was Xeno™ medium containing trypsin (20 U/mL) to maintain a constant trypsin activity in the STR to ensure cleavage of the IAV hemagglutinin protein to facilitate virus entry.

### Sampling of the STR

Samples were taken with a syringe via a sample port. Sampling was possible either from the cultivation vessel or from the tubing behind the cell retention membrane (permeate line). This allowed to measure and calculate the percentage of PFU and DI244 vRNA that passed through the cell retention membrane (Eq. 4) for each individual sampling time point. Part of the sample was used for VCC measurements (Vi-CELL XR, Beckman Coulter, 731050), the rest was centrifuged (3000g, 10 min, 4°C) and the supernatant was aliquoted and stored at −80°C for subsequent analysis in the interference assay, virus quantification assays or metabolite analysis (Bioprofile 100 plus analyzer, Nova Biomedical).

### Interference assay

The interfering efficacy of the produced DI224 was evaluated by a previously established assay (Hein et al. 2021a; Hein et al. 2021b). Briefly, MDCK cells were co-infected with STV at MOI 10 or 0.01 and 125 μL of the produced DI244. As control, infections with “STV only” were carried out. Comparison of co-infections and the control infection allowed an assessment of the DI244-induced reduction of released virus particles. DI244 preparations resulting in a more pronounced titer reduction were considered to have a higher interfering efficacy. Addition of DI244 material with a fixed volume allowed evaluation of the interfering efficacy per product volume and therefore the identification of optimal production conditions.

### Virus quantification assays

The total amount of IAV particles was quantified using a hemagglutination assay and was expressed as log_10_ hemagglutinin units/100 μL (log_10_ HA units/100 μL) (Kalbfuss et al. 2008). PFU were determined by a plaque assay, conducted as described previously (Hein et al. 2021a; Hein et al. 2021b; Kupke et al. 2020). Here, for the quantification of released infectious virus particles in the interference assay, parental adherent MDCK cells were used, only allowing propagation of STVs. For quantification of DI244 in pure DIP preparations, MDCK-PB2(adh) cells were used. The virus titer or DI244 titer was expressed as PFU/mL. The relative standard deviation (RSD) of technical duplicates was ≤ 43.8%.

### Real-time reverse transcription qPCR

For quantification of genomic vRNA, a real-time reverse transcription qPCR (real-time RT-qPCR) was used. The vRNA in the cell culture supernatant was purified using a NucleoSpin^®^ RNA virus kit (Macherey-Nagel, 740956) according to manufacturer’s instructions. The real-time RT-qPCR was conducted according to a previously reported method (Kawakami et al. 2011). This method allows for a gene-specific quantification of individual IAV vRNA segments. The adaptations made to the protocol as well as the procedure for standard generation, RT, real-time PCR and absolute quantification of vRNA levels have been reported previously (Frensing et al. 2016; Kupke et al. 2019). For quantification of the shortened DI244 vRNA, an assay using specific primers binding across the junction region of the deletion was used (Wasik et al. 2018). The RSD of technical quadruplicates was ≤ 52.5%.

### Protein and DNA analysis

Before the analysis of protein and DNA content in the produced material, samples were centrifuged (3000g, 10 min, 4°C) and stepwise micro-filtered (0.45 μm filtration followed by a 0.2 μm filtration). Protein content in produced DI244 material was determined by a Bradford assay (BioRad Laboratories, 500-0006). For the standard calibration curve bovine serum albumin (Sigma-Aldrich Chemie GmbH, A3912) was used. Double-stranded host cell DNA (dsDNA) was quantified by a Quant-iT PicoGreen assay (Life Technologies GmbH, P7581) as described previously (Opitz et al. 2007). The standard calibration curve was made using lambda DNA (Promega, D1501).

### Calculations

The total amount of produced DI244 particles for each cultivation was calculated using the plaque titers in the cultivation vessel and in the permeate bottle at time of harvest.

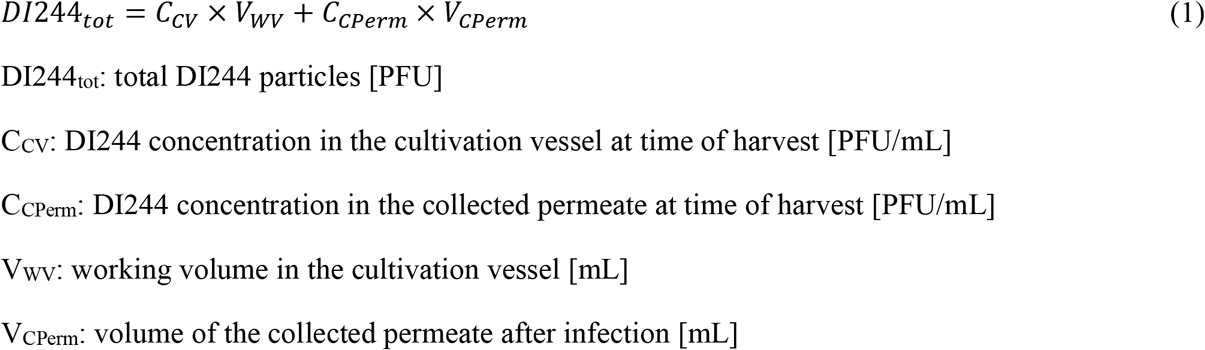

Based on the total amount of produced DI244 particles the PFU per cell were calculated as followed.

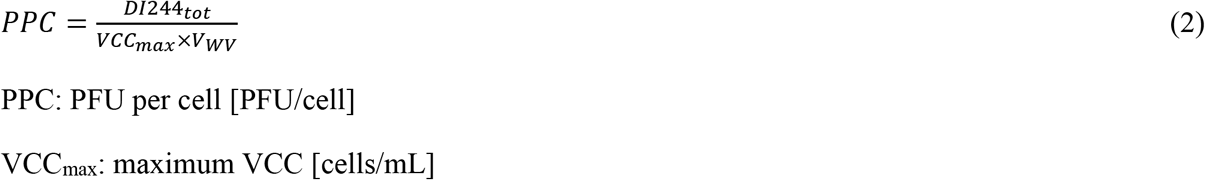

Perfusion with the VHU allows for harvesting of DI244 particles through the membrane. Here, virus titer in the cultivation vessel and in the collected permeate differed at time of harvest. To simplify the reporting of virus titers, the theoretical overall virus titer of the entire virus containing harvest was calculated.

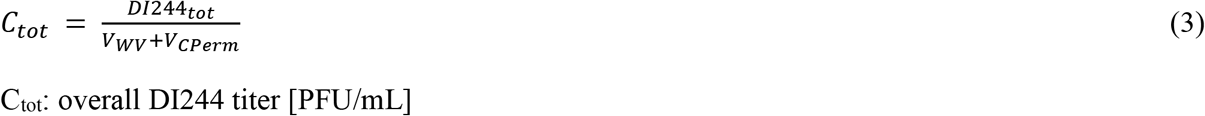

The DI244 titer in the cultivation vessel and in the permeate line were determined for every sample time point. This allowed the calculation of the percentage of PFU passing through the membrane. The overall percentage of passed PFU was calculated as the average of all sampling time points.

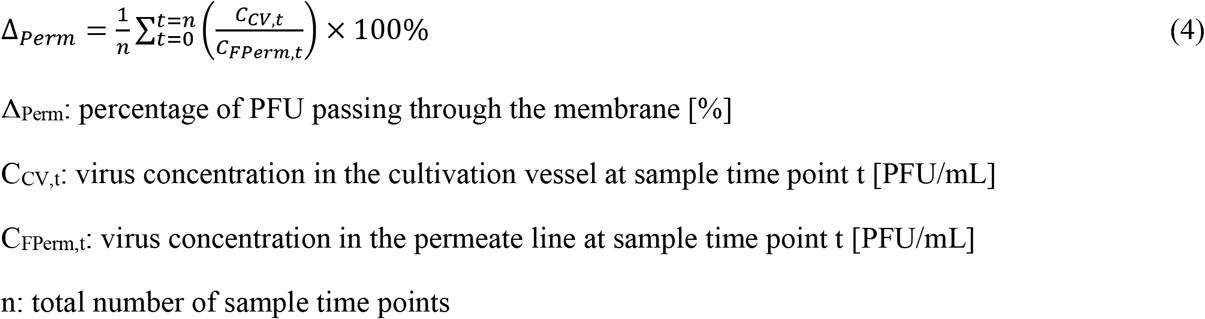

Please note that Eq. 1 – 4 can be easily modified to calculate the total RNA copies (Eq. 1), the vRNA copies per cell (Eq. 2), the overall vRNA level (Eq. 3), and the percentage of DI244 vRNA copies passing through the membrane (Eq. 4). Here, the measured vRNA level must be used instead of the plaque titer.

## Results

### Manually adjusted and automated perfusion cultivations resulted in high cell concentrations and IAV DIP titers

We previously reported the batch production of pure DI244 particles in shake flasks for the use as an antiviral (Hein et al. 2021a). To evaluate if process intensification strategies that are already used for non-modified MDCK suspension cells and IAV STV production (Wu et al. 2021) would be applicable, a manually adjusted perfusion process in a STR was implemented. For this cultivation, the perfusion rate was adjusted every 12 h, according to a pre-calculated profile. The profile was based on the cell growth rate and metabolite uptake rates observed in a previous batch cultivation (Fig. S1) (Hein et al. 2021a).

In a next step, the perfusion rate was controlled using a capacitance probe. Here, the signal was correlated to the VCV in the STR. For the manually adjusted perfusion cultivation, an average CSPR of 200 pL/cell/d was realized, for the controlled perfusion cultivations a CVSPR of 0.12 pL/μm^3^/d was chosen. This would equal a CSPR of 200 pL/cell/d for cells with a diameter of 15 μm. One manually adjusted and three automated perfusion cultivations were performed in the STR. The manually adjusted and one automated perfusion cultivation were conducted with a commonly used HFM to compare against results obtained with non-modified MDCK suspension cells and IAV STR production (Wu et al. 2021). For the following two automated perfusion cultivations, the VHU was evaluated as a new cell retention membrane to test options for direct harvesting of virus particles through the membrane.

Similar cell growth was observed for both perfusion strategies and both cell retention membranes reaching cell concentrations higher than 20E+6 cells/mL with viabilities above 95% (Fig. 2a). Further, no membrane fouling or blocking of either membrane was observed, indicating that both membranes are equally suited for high cell density cultivations.

**Fig. 2.**
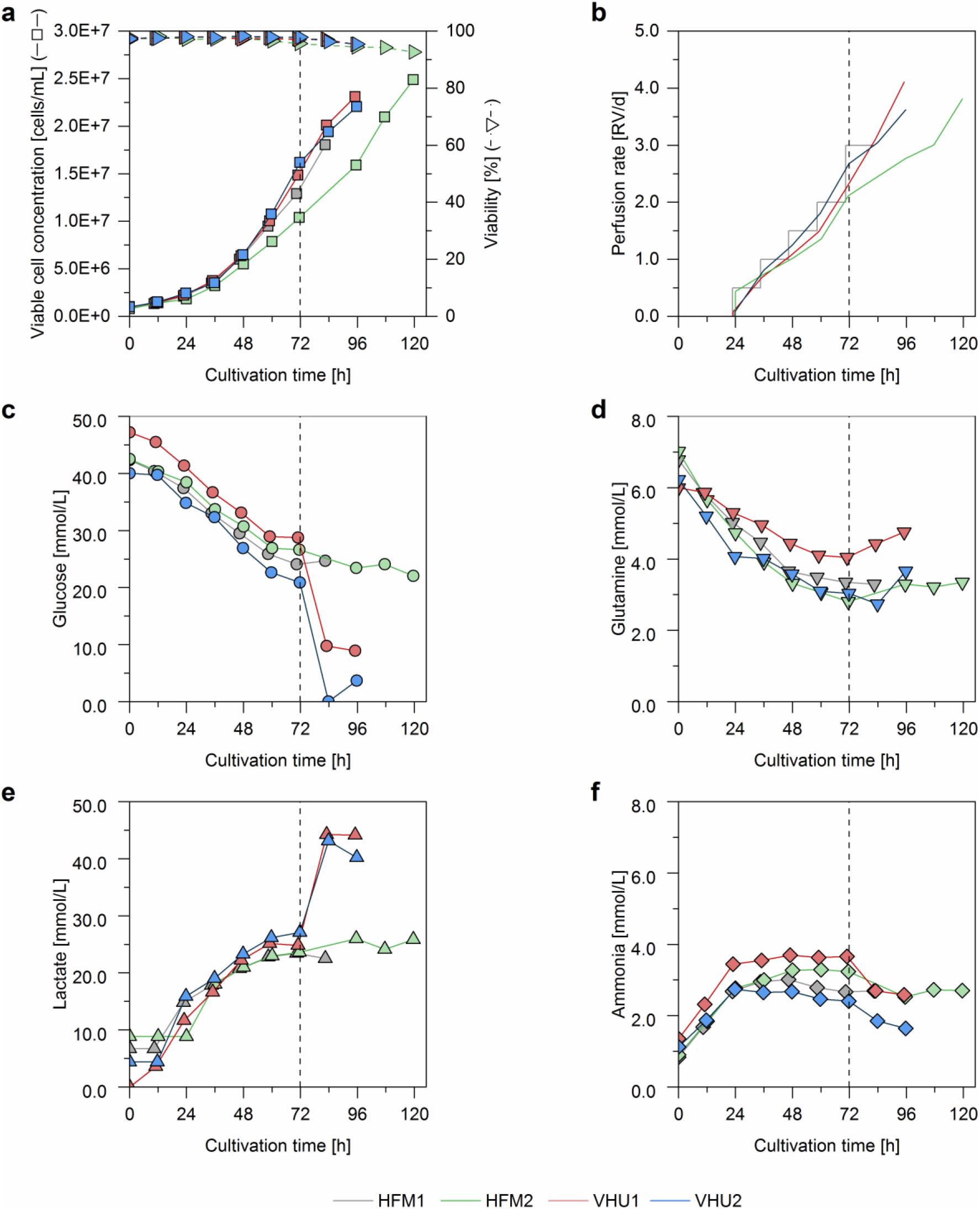
Cell growth and metabolite concentrations of MDCK-PB2(sus) cells cultivated in a 1 L stirred tank bioreactor coupled to an alternating tangential flow filtration system (ATF 2). The first perfusion cultivation was adjusted manually (HFM1), the remaining cultivations were controlled based on cell volume monitoring using a capacitance probe (Fig. S2). For the automated perfusion cultivations, gassing with an additional micro-sparger was initiated 72 hours post inoculation (indicated by vertical dashed line). As cell retention device a commonly used hollow fiber membrane (pore size of 0.2 μm; HFM1 and HFM2) or the virus harvest unit (pore size of ~10 μm; VHU1 and VHU2) was used. (a) Viable cell concentration and viability, (b) perfusion rate (determined by the permeate weight), (c) glucose concentration, (d) glutamine concentration, (e) lactate concentration, (f) ammonia concentration.

The automated perfusion control was successfully implemented. The permittivity signal showed a linear correlation with the VCV in the STR (Fig. S2) and the perfusion rate automatically increased with increasing VCV (Fig. 2a–b). For all four perfusion cultivations, the metabolite levels were very comparable and no substrate limitation or inhibition by waste products was observed until 72 h post inoculation (Fig. 2c–f). For the manually adjusted perfusion cultivation (HFM1), oxygen limitations were observed 84 h post inoculation (data not shown). To circumvent this in the subsequent automated perfusion cultivations, gassing with an additional micro-sparger was initiated 72 h post inoculation. This did not seem to have an impact on the cell metabolism when the HFM was used (Fig. 2c – f). Nevertheless, this cultivation showed a slightly lower overall growth rate perfectly reflected by the perfusion rates of the automated control. When the VHU was used, the additional sparging seemed to result in increased glucose uptake and lactate production together with a slight ammonia uptake instead of a release (Fig. 2c, e, f). Nevertheless, at this point, differences in cell growth compared to the manually adjusted cultivation seem to be negligible. Overall, for all four cultivations, up to 20E+6 cells/mL were achieved for subsequent DI244 production.

### Perfusion cultivations with the VHU allowed continuous harvesting of influenza A virus DIPs

Next, the cell retention membranes were compared for their ability to allow for harvesting virus particles into the permeate. Here, cells were infected when the VCC reached concentrations above 20.0E+06 cells/mL (or 25.0E+06 cells/mL for HFM2). Maintaining a constant pH value for cultivation HFM2 at cell concentrations up to 26.8E+6 cells/mL required addition of large amounts of base (7.5% NaHCO_3_ solution). Therefore, it was decided to infect subsequent cultivations (VHU1 and VHU2) before the VCC surpassed 25.0E+6 cells/mL. Before infection of cells with a pure DI244 seed virus at MODIP 1E–3, the medium was fully exchanged, the temperature reduced from 37°C to 32°C and trypsin added to 20 U/mL. The permeate flow was paused for one hour post infection to avoid loss of trypsin and virus particles into the permeate and to allow for an efficient cell entry of the DIPs. During the virus replication phase, cultivation medium containing 20 U/mL trypsin was used and the perfusion rate was kept constant at 2–3 RV/d.

For all four perfusions, cells continued to grow for the first 6–12 hours post infection (hpi) and reached VCC up to 28.4E+6 cells/mL (Fig. 3a). Afterwards, viability and VCC started to decline. HA titers (Fig. 3b), vRNA levels (Fig. 3c), and PFU titers (Fig. 3d) all reached their respective maximum at about 36 hpi. The maximum plaque titer was 7.4E+9 PFU/mL and the maximum DI244 vRNA level was 5.9E+11 copies/mL, both being more than 10-times higher than for a batch production process infected at a VCC of 2.0E+06 cells/mL described before (Hein et al. 2021a). In contrast to previous findings (Genzel et al. 2010; Hein et al. 2021a), no decrease in the plaque titer was observed after the respective maximum was reached. The increased virus stability could be explained by the reduced cultivation temperature of 32°C during the virus replication phase and was confirmed by small scale experiments (Fig. S3).

**Fig. 3.**
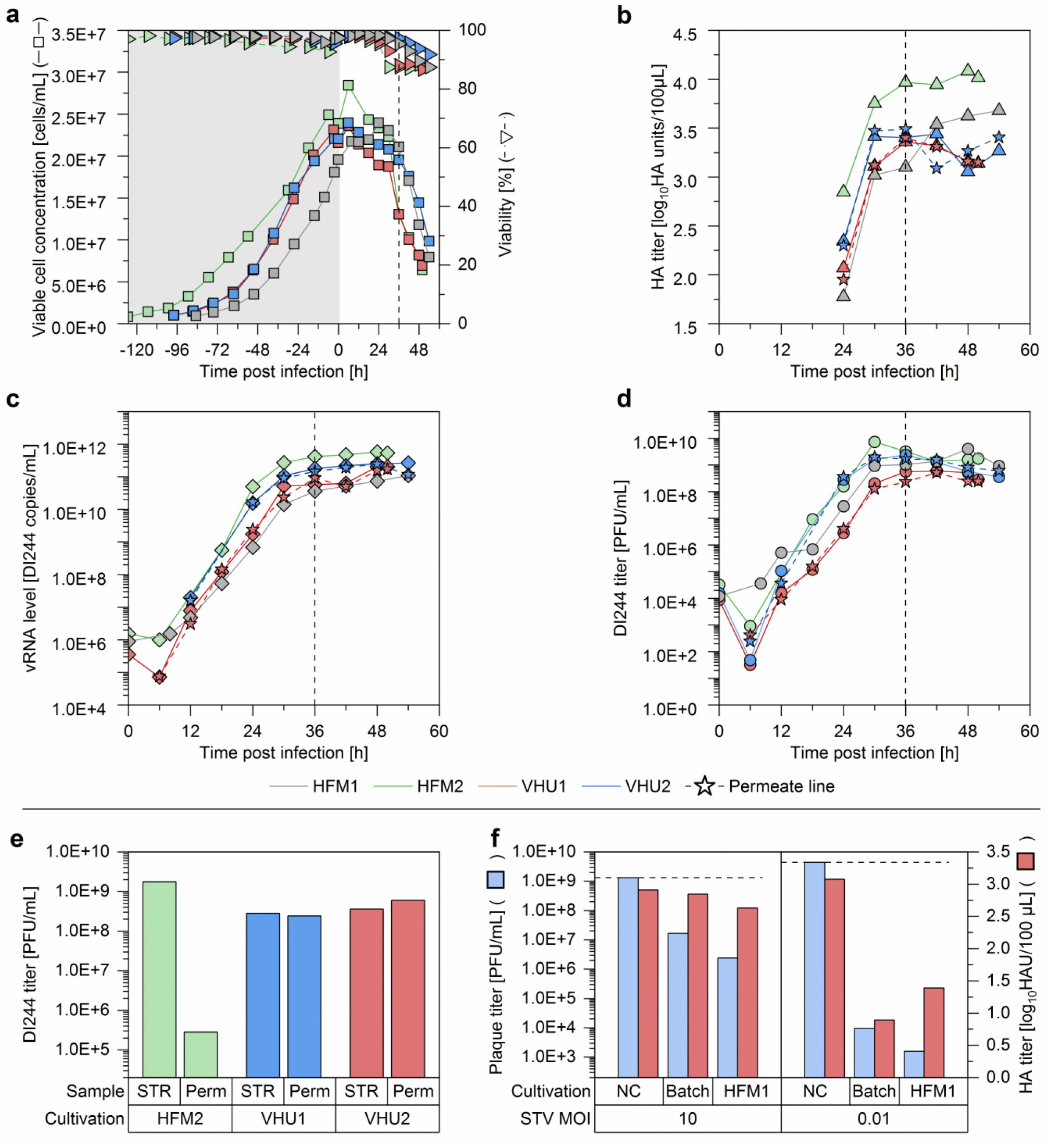
Influenza A virus DIP production with MDCK-PB2(sus) cells cultivated in a 1 L stirred tank bioreactor coupled to an alternating tangential flow filtration system (ATF 2). The perfusion rate of one cultivation was adjusted manually (HFM1), the remaining cultivations were controlled using a capacitance probe for cell growth monitoring. As cell retention devices a commonly used hollow fiber membrane (pore size of 0.2 μm; HFM1 and HFM2) or a virus harvest unit (pore size of ~10 μm; VHU1 and VHU2) was used. MDCK-PB2(sus) cells were infected at a concentration of 25.0E+6 cells/mL (HFM2) or 20.0E+6 cells/mL (HFM1, VHU1, VHU2) with a pure DI244 seed virus at MODIP 1E–3. (a) Viable cell concentration and viability (grey area indicates cell growth phase), (b) HA titer, (c) DI244 vRNA level, (d) DI244 titer. (b, c, and d) solid lines indicate virus titer in the cultivation vessel, dashed lines with star symbols indicates the virus titer in the permeate line. Vertical dashes line indicates time when maximum titers were reached (36 hpi). (e) DI244 titer in cultivation vessel (STR) and permeate line (Perm) at time of harvest (48 hpi). (f) Interference assay for DI244 produced in HFM1. For comparison, previously reported DI244 material (Batch; produced in shake flask with infection at 2.0E+6 cells/mL and MODIP 1E–2) (Hein et al. 2021a) or medium as negative control (NC) was tested.

Additionally to the virus titers of the cultivation vessel, titers in the permeate line were determined (Fig. 3b–d). For the used HFM it is well established that virus particles of about 40-100 nm in diameter cannot pass through the membrane (Nikolay et al. 2020). Samples taken shortly before harvesting confirmed this, since only residual virus amounts could be detected in the permeate line (Fig. 3e). However when the VHU was used, the virus titer in the cultivation vessel always equaled the virus titer in the permeate line (Fig. 3b–e). Here, the percentage of PFU and DI244 vRNA passing through the membrane was calculated as about 100% for each sampling time point (Eq. 4; Tab. 2). Both values were more or less stable over the cultivation time, which indicates that no filter fouling or blocking occurred even at later process times where high amounts of cell debris were present in the culture broth. This clearly shows the potential of the VHU for continuous virus harvesting during a perfusion cultivation.

**Table 2.**
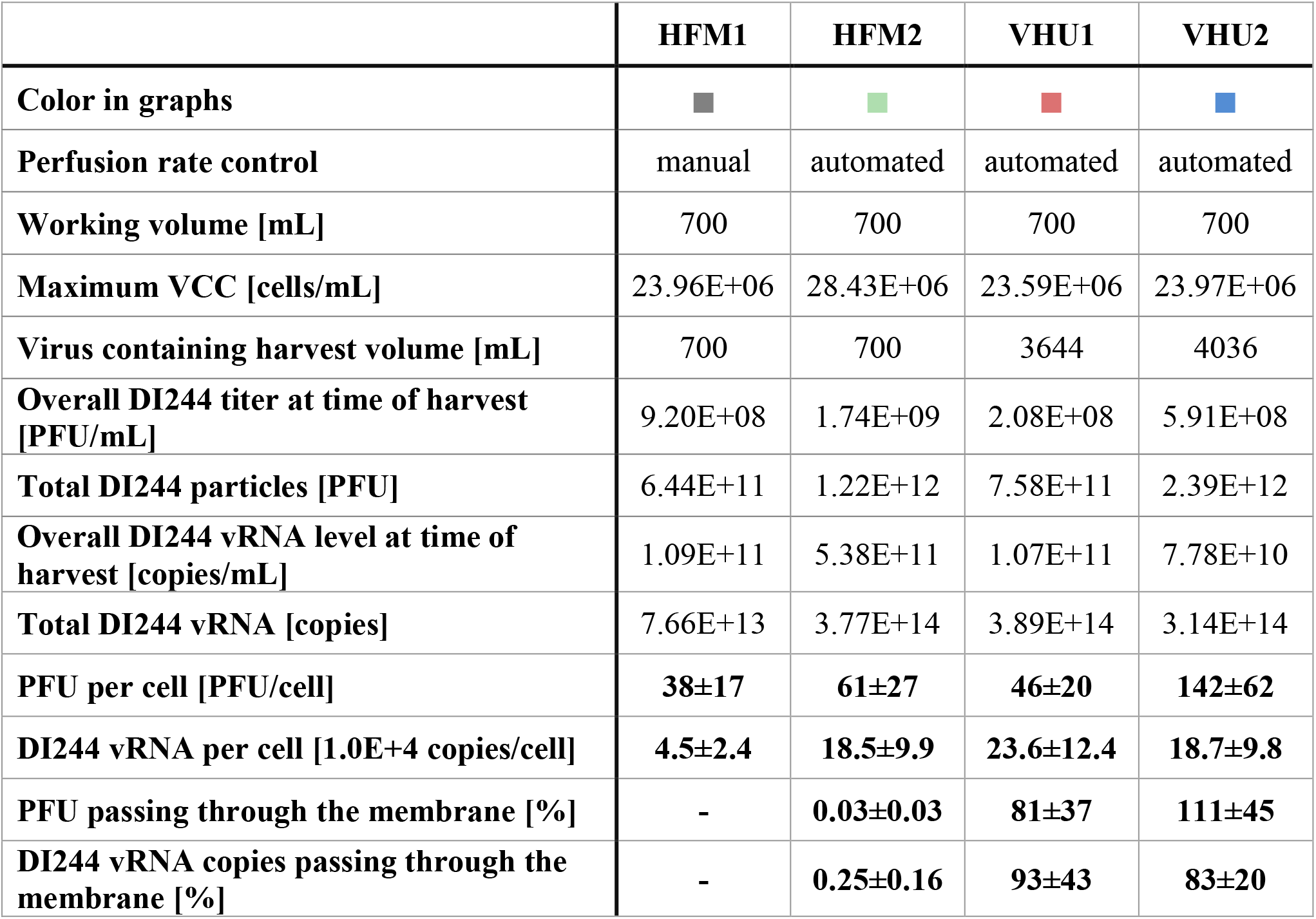
Overview of process parameters, virus titers and calculated yields for influenza A virus DIPs produced in a 1 L stirred tank bioreactor coupled to an alternating tangential flow filtration system (ATF 2). Perfusion rates for cultivations with MDCK-PB2(sus) cells were either adjusted manually or controlled using a capacitance probe for cell growth monitoring. As cell retention device a commonly used hollow fiber membrane (pore size of 0.2 μm; HFM1 and HFM2) or a virus harvest unit (pore size of ~10 μm; VHU1 and VHU2) was used. All cultures were harvested at 48 hpi. For the ratio PFU/vRNA per cell, the error based on the relative standard deviation of the assay is given. For the percentage of PFU/vRNA passing through the membrane the average of the whole virus replication phase is given with the corresponding standard deviation. Percentages of PFU/vRNA passing through the membrane higher than 100% were calculated when the measured titer in the permeate line was higher than in the STR at the same sample point. This should theoretically not be possible and can be attributed to the high standard deviation of virus quantification assays.

For perfusion cultivations with the VHU, virus particles were diluted as they also passed on to the permeate. Therefore, the achieved virus titers were lower (Fig. 3b–d). For a more detailed comparison of the tested cell retention membranes, the overall process yield, the total amount of virus particles, and the virus yields were calculated (Tab. 2). The total amount of DI244 particles, the total amount of DI244 vRNA, PFU per cell, and the vRNA per cell were similar using either membrane. Although the average values of perfusion cultivations using the VHU were a bit higher, the differences were not significant, given the relatively high standard deviation of the virus quantification assays.

Finally, DI244 material produced in a perfusion process (HFM1) and harvested at 48 hpi was tested in an interference assay to verify its antiviral potential compared to material produced previously in a batch cultivation (Hein et al. 2021a) (Fig. 3f). In the interference assay, co-infections of STV (MOI 10 or 0.01) and produced DIP material (fixed volume) were carried out. DIP material resulting in a more pronounced titer reduction is considered to have a higher interfering potency. As the DI244 titer was more than 10-times higher for material produced in the perfusion cultivation, a higher interfering potency was expected. Indeed, the DI244 material produced in a perfusion cultivation reduced the plaque titer by almost one order of magnitude more than material produced in a batch cultivation. For the HA titer a similar trend could be observed when STV was added at a MOI of 10, but not for STV MOI 0.01. Here, it should be considered that DI244 particles themselves express HA and contribute to the HA titer. Overall, the results of the interference assay clearly demonstrate that DI244 produced in a perfusion cultivation has a high interfering efficacy and should be suited for antiviral therapy.

### Retention of proteins and dsDNA depends on the cell retention device

Continuous harvesting of virus particles during a perfusion process theoretically allows the implementation of process integration strategies, where the harvested material can be directly transferred into subsequent downstream processing units without hold times. However, besides virus particles, the cultivation broth contains intact cells, cell debris, and host cell proteins as well as host cell DNA released with cell lysis. The amount of these contaminations that are able to pass through the membrane could have severe implications regarding the performance of downstream processing. Therefore, the protein and dsDNA contamination level in the produced virus material was determined.

For the perfusion with the HFM2, only the cultivation vessel contained virus particles and was harvested (700 mL). For the perfusion with the VHU, the permeate also contained virus particles and was harvested additionally. Consequentially, the harvest volume was much higher (3644 mL). First, the protein and dsDNA concentration in the virus containing harvest was measured. The VHU appeared to not retain any contaminations, as the protein and dsDNA concentration in the cultivation vessel and in the permeate were more or less the same (Fig. 4a). Since proteins and dsDNA were heavily diluted over the time of the cultivation when the VHU was used, their contamination level was very low. In strong contrast, the HFM withhold most proteins and dsDNA and much higher levels were observed. The drastically reduced host cell protein retention of the VHU might also indicate advantages over the HFM for continuous harvesting of recombinant proteins.

**Fig. 4.**
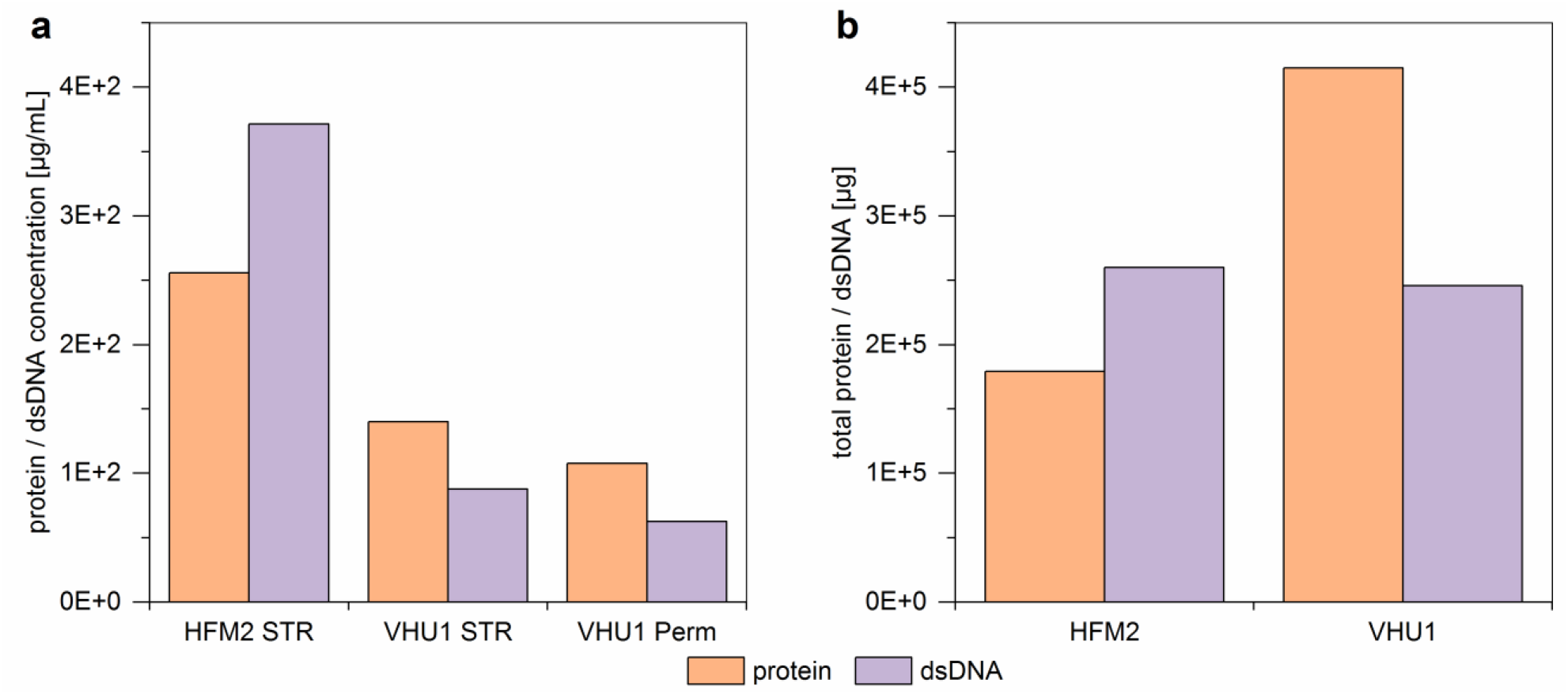
Protein and dsDNA contamination in the virus containing harvest of influenza A virus DIPs produced in 1 L stirred tank bioreactors coupled to an alternating tangential flow filtration system (ATF 2). The harvest (48 hpi) of two perfusion cultivations using either a commonly used hollow fiber membrane (pore size of 0.2 μm; HFM2) or a virus harvest unit (pore size of ~10 μm; VHU1) were analyzed for their (a) protein and dsDNA concentration. For HFM2, only the cultivation vessel (700 mL) was harvested; for VHU1, the cultivation vessel (STR; 700 mL) and the permeate (Perm; 2944 mL) were harvested. (b) total amount of protein and dsDNA based on the respective harvest volume.

Considering the respective harvest volumes, the total amount of proteins and dsDNA were calculated (Fig. 4b). The HFM appeared to retain the entire dsDNA, since the total amount of dsDNA found in the cultivation vessel equaled the amount determined for the VHU in the total virus harvest volume (cultivation vessel and collected permeate). In contrast, the HFM appeared to only partly withhold proteins and the total amount of proteins contaminating the virus harvest was much lower.

## Discussion

### MDCK-PB2(sus) cells can be used for high cell density cultivations

The metabolite uptake rates of MDCK-PB2(sus) cells grown in perfusion cultivations were very comparable to those reported previously for batch cultivations and to data obtained for the parental suspension cell line (descendent from a MDCK cell line originally obtained from ECACC) (Hein et al. 2021a). Therefore, expression of PB2 itself did not seem to significantly influence cell metabolism or cell growth. However, uptake rates are relatively high and consequentially a high CSPR of 200 pL/cell/day was needed for perfusion cultivations. For comparison, when a perfusion process with the same medium but a different MDCK cell line (descendent from a MDCK cell line originally obtained from ATCC) was established, a lower CSPR from 60 pL/cell/day was sufficient (Wu et al. 2021). Furthermore, no oxygen limitations were observed in cultivations with the latter and the optimal pH value for cell growth was much lower (pH 7.5 for MDCK-PB2(sus) and its parental cell line (ECACC); pH 7.0 for MDCK cells obtained from ATCC). Previous studies compared the cell growth and virus yield of MDCK cells obtained either from ECACC or ATCC grown in Xeno™ medium (Bissinger 2020). Here, faster cell growth, higher maximum VCC and a higher PFU per cell were observed for MDCK cells obtained from ATCC. Therefore, for further improving process performance, it would be an option to genetically modify MDCK cells from ATCC to express the viral PB2 and evaluate growth and productivity at high cell concentrations.

Nevertheless, perfusion control using a capacitance probe for cell growth monitoring was sufficient to maintain high substrate and low waste product levels over the cultivation time. The automation could allow for a robust process control for even higher cell concentrations, as already shown for other cell lines (Nikolay et al. 2018). In the present study, a maximum concentration of 28.4E+06 cells/mL was reached. Here, it should be possible to achieve even higher VCC with further optimization of the pH and DO control. With the current control regime both parameters were hard to maintain at high cell concentrations and it was decided for subsequent cultivations to infect the cells, before the VCC exceeded 25.0E+6 cells/mL. To overcome oxygen limitations observed for the manual perfusion cultivation, aeration with an additional micro-sparger was started 72 hours post inoculation for the automated perfusion cultivations. Here, increased glucose uptake and lactate production was observed, when the VHU was used. The increased lactate formation might indicate increased cell stress. For example, it was reported that cells exposed to shear stress show an increased lactate dehydrogenase activity (Shiragami and Unno 1994). One possible explanation for the observed change in metabolism could be that the tubular VHU membrane caused higher shear stress than the HFM, in particular, when combined with a micro-sparger. This could have implications on the maximum achievable VCC and will be further investigated in future studies in our lab.

In summary, MDCK-PB2(sus) cells showed great potential for the design of high cell density processes to achieve very high DI244 titers. Further improvements of the cell line and process control could allow for even higher cell concentrations and virus titers.

### Virus harvest units allow for continuous virus harvesting

Previous studies investigated different HFMs for their potential to allow continuous virus harvesting (Genzel et al. 2014; Nikolay et al. 2020). However, almost all tested membranes completely retained the produced virus particles, despite the nominal pore size reported was much larger than the virus diameter. It was concluded that the membrane material and with this, its inner surface and structure, porosity, and hollow fiber wall thickness largely influence the retention of virus particles. In the present study, these findings were confirmed for a commonly used HFM (nominal pore size 0.2 μm) that retained most IAV DIPs (approximate diameter 80–120 nm). In strong contrast, the use of the VHU allowed approximately 100% of the produced IAV DIPs to pass through the membrane into the permeate. To our knowledge, this is the first time that continuous virus harvesting through a membrane with a high yield was demonstrated for perfusion cultivations. This is especially remarkable since IAV is a lytic virus and, therefore, large amounts of cell debris accumulate towards the end of cultivations. In particular, no filter fouling or blocking was observed as the percentage of PFU and DI244 vRNA passing through the membrane remained unchanged over the entire virus production phase. Furthermore, the VHU did not retain host cell proteins in contrast to the used HFM. This might also indicate an advantage over conventional HFM for production and harvest of recombinant proteins (e.g. monoclonal antibodies).

### Continuous virus harvesting with non-membrane-based perfusion systems compared to the membrane-based perfusion cultivation using the VHU

Besides membrane-based cell retention devices there also exist other systems. The potential of two of these systems (acoustic settler and inclined settler) for continuous virus harvesting was described recently for IAV and MVA production (Coronel et al. 2020; Granicher et al. 2020; Granicher et al. 2021). Choosing one of these systems will depend on many parameters and a detailed comparison of pros and cons can be found elsewhere (Chotteau 2015). In our case, the operation of an ATF system seems to have advantages as system cooling or complex pumping strategies are not required. Further, it was described that the mentioned alternative cell retention systems do not allow for complete cell retention and some cells are lost over the course of cultivations. Additionally, their separation principles rely on cell settling and therefore longer recirculation times were observed (Chotteau 2015; Coronel et al. 2020; Granicher et al. 2020). In the present study, no cells could be detected in the permeate line of the VHU (data not shown) and the cells only spent a few seconds outside the STR during each pumping cycle of the ATF system. Finally, both the acoustic and the inclined settler do not offer the possibility to retain virus particles. With the ATF system it is now possible to retain or continuously harvest the produced virus particles depending on the membrane chosen for cultivation. In summary, the VHU allows the set-up of a continuous virus harvesting strategy in already established ATF perfusion cultivations only by substituting the used cell retention membrane.

### Influenza A virus DIPs harvested from a perfusion process show potential for future production of antivirals

The potential of IAV DIPs, and DI244 in particular, as an antiviral was already shown *in vivo* (Dimmock et al. 2008). Yet, the material investigated in those studies was produced in eggs. We recently also demonstrated the antiviral potential of purely clonal cell culture-based produced DI244 *in vivo* (Hein et al. 2021a). Here, 100% of mice infected with an otherwise lethal IAV infection survived when treated with the produced DI244. However, relatively large amounts of DI244 (1.5E+6 PFU per mouse) were administered. Therefore, it was important to establish a production process that could supply the needed quantities. The average amount of produced DI244 for the four perfusion cultivations with a working volume of 700 mL was 1.25E+12 PFU. This would be enough to treat millions of mice. While the dose for humans is currently unknown, it seems very likely that even a small scale perfusion process could be used to generate a high number of DI244 doses for human use. Overall, this provides further support for the feasibility of cell culture-based production of DIPs for antiviral treatment.

## Conclusion

In this study, we established an automated perfusion process for the production of purely clonal IAV DIPs using a cell culture-based production platform. To our knowledge, this is the first time pure DIP preparations have been produced in perfusion mode. The very high DIP titers obtained clearly demonstrate the feasibility of cell culture-based DIP production for antiviral therapy. Using a VHU as a membrane-based cell retention device allowed continuous virus harvesting during the perfusion cultivation. As far as we are aware, this is the first time that this process option was described. It opens up new possibilities for process integration not only for virus production but equally for recombinant protein production in perfusion mode.

## Declarations

### Ethics approval

Not applicable.

### Consent to participate

Not applicable.

### Consent for publication

Not applicable.

### Availability of data and material

The datasets generated and/or analyzed during the current study, cell lines, and seed virus are available from the corresponding author on reasonable request.

### Author Contributions

Conceptualization, MH, AC, MC, YG, and UR; Methodology, MH and YG; Investigation, MH and AC; Writing – Original Draft, MH; Writing – Review & Editing, MH, AC, MC, SK, YG and UR; Supervision, YG, SK and UR; Project Administration, MH and YG; Funding Acquisition, UR.

## Acknowledgments

The authors would like to thank Nancy Wynserski and Claudia Best for excellent technical assistance and Pavel Marichal-Gallardo for support with protein and DNA analysis. The authors greatly appreciate the contribution of Shanghai BioEngine Sci-Tech and Prof. Tan from the East China University of Science and Technology for providing the Xeno™ medium. The authors would like to thank the German Primate Center-Leibniz Institute for Primate Research and Prof. Pöhlmann for generating and providing the MDCK-PB2(sus) cell line.

