## Supplementary file 1 for "Cell culture-based production of defective interfering influenza A virus particles in perfusion mode using an alternating tangential flow filtration system"

Journal: “Applied Microbiology and Biotechnology”

### Supplementary Material

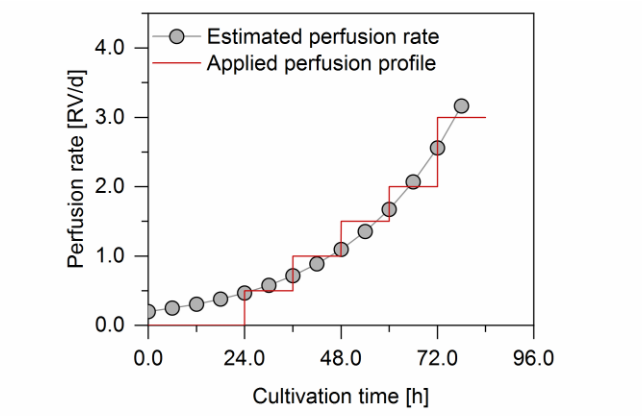


**Fig. S1 Estimated and manually adjusted perfusion rate for cultivation of MDCK-PB2(sus) cells in a 1 L stirred tank bioreactor (HFM1).** For the calculation of the required perfusion rate, the maximum specific growth rate (0.0354 1/h) and the glucose uptake rate (3.62E–10 mmol/cell/h) of a previous cultivation was used. Considering the metabolite uptake rates and the glucose (40 mmol/L) and glutamine (8 mmol/L) concentration in the Xeno™ medium, a cell specific perfusion rate (CSPR) of 200 pL/cell/d was calculated. Based on the expected cell growth and the CSPR the required perfusion rate was estimated. A stepwise perfusion profile was chosen accordingly.

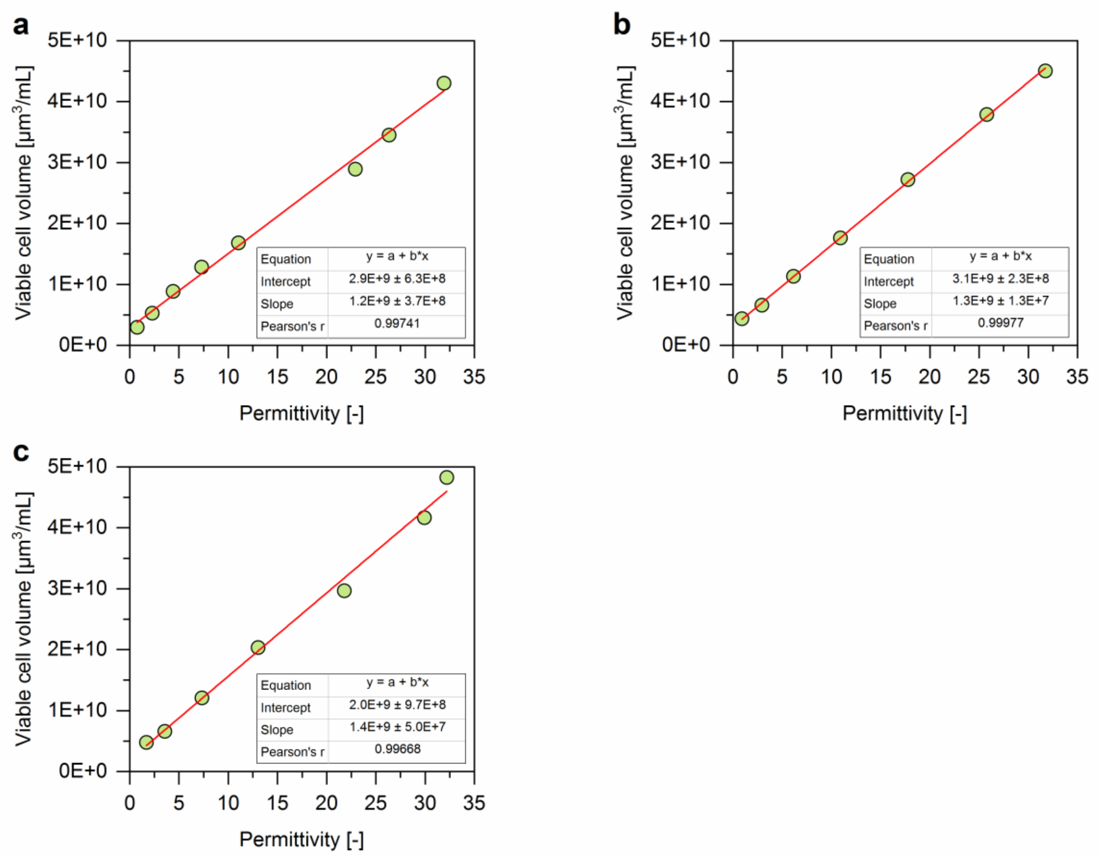


**Fig. S2 Linear regression of the viable cell volume and permittivity measured for MDCK-PB2(sus) cells cultivated in a 1 L stirred tank bioreactor coupled to an alternating tangential flow filtration system (ATF 2).** Perfusion cultivations were controlled using the capacitance probe for cell growth monitoring. As cell retention device a commonly used hollow fiber membrane (pore size of 0.2 µm; HFM2) or the virus harvest unit (pore size of ~10 µm; VHU1 and VHU2) was used. (a) HFM2, (b) VHU1, (c) VHU2.

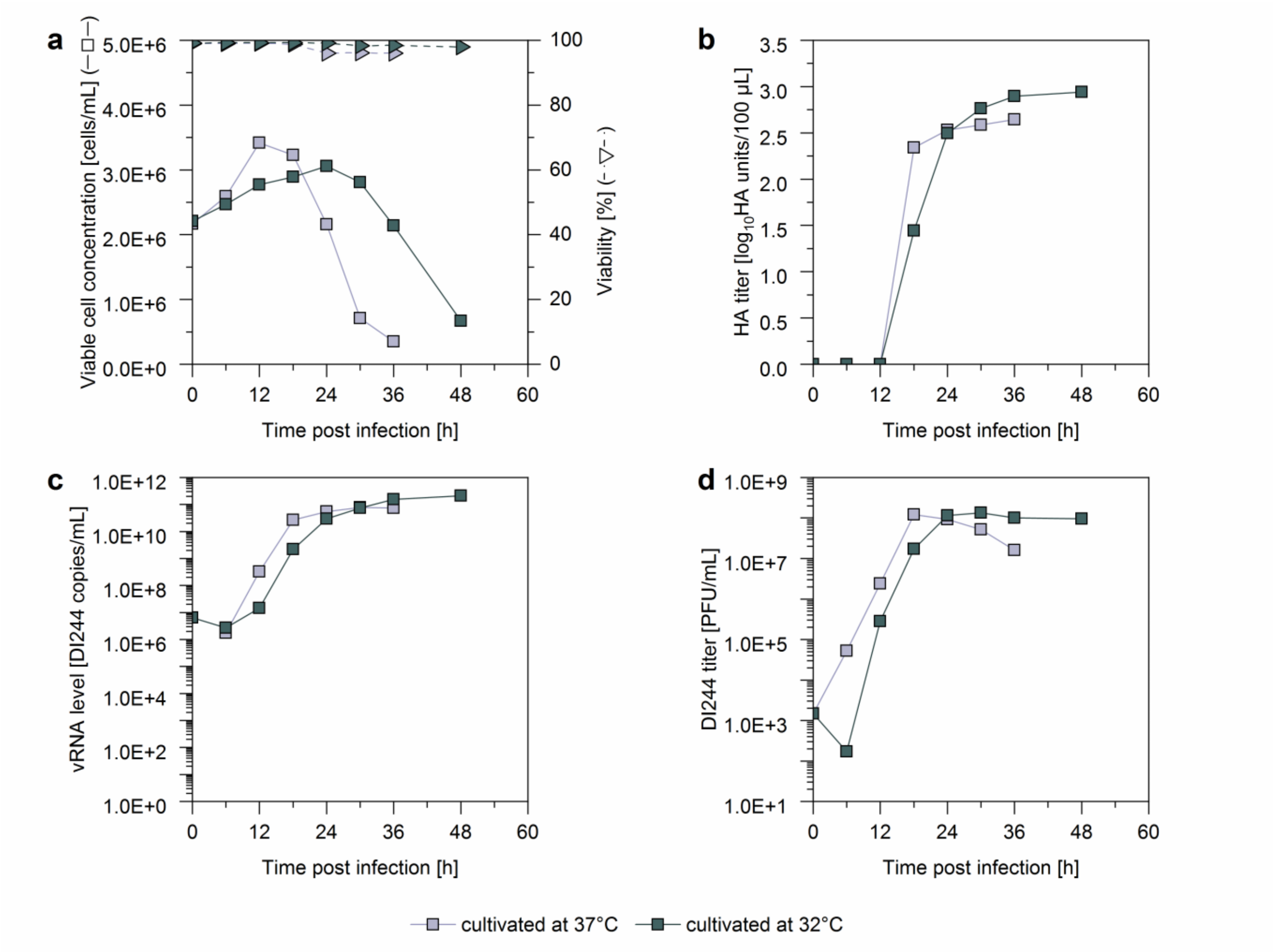


**Fig. S3 Influenza A virus DIP production with MDCK-PB2(sus) cells cultivated in a shake flask (40 mL working volume) at different temperatures.** At time of infection the viable cell concentration (VCC) was adjusted to 2.0E+6 cells/mL, the cells were infected with a pure DI244 seed virus at MODIP 1E–3. After infection, the cells were split and either cultivated at 37°C or 32°C. (a) VCC and viability, (b) HA titer, (c) vRNA level, and (d) DI244 titer.
